# Oxygen-induced chromophore degradation in the photoswitchable red fluorescent protein rsCherry

**DOI:** 10.1101/2023.01.13.523900

**Authors:** Thi Yen Hang Bui, Elke De Zitter, Benjamien Moeyaert, Ludovic Pecqueur, Bindu Y Srinivasu, Anastassios Economou, Marc Fontecave, Luc Van Meervelt, Peter Dedecker, Brandán Pedre

## Abstract

Reversibly switchable monomeric Cherry (rsCherry) is a photoswitchable variant of the red fluorescent protein mCherry. We report that this protein gradually and irreversibly loses its red fluorescence in the dark over a period of months at 4°C and a few days at 37°C. We also find that its ancestor, mCherry, undergoes a similar fluorescence loss but at a slower rate. X-ray crystallography and mass spectrometry reveal that this is caused by the cleavage of the *p*-hydroxyphenyl ring from the chromophore and the formation of two novel types of cyclic structures at the remaining chromophore moiety. Overall, our work sheds light on a new process occurring within fluorescent proteins, further adding to the chemical diversity and versatility of these molecules.

## 1. Introduction

Fluorescent proteins (FPs) are amongst the most ubiquitously used protein tags in life science research, providing the ability to genetically label proteins with intrinsically fluorescent indicators [1]. Of particular interest are reversibly photoswitchable fluorescent proteins (RSFPs). RSFPs typically exist in a bright fluorescent state and can be reversibly and controllably switched between a fluorescent (on) and a non-fluorescent (off) state using light of specific wavelengths. This photoswitching can be repeated up to hundreds of times and forms the basis of super-resolution imaging techniques such as RESOLFT or pcSOFI [2].

Most of the available RSFPs emit green fluorescence and are characterized by a high molecular brightness, high switching contrast, and a high fatigue resistance [3–6]. RSFPs with other emission colors are also available, including cyan, yellow/orange, and red fluorescent proteins [7–10]. While the red-emitting RSFPs have advantages for low-background or even in-tissue imaging, their optical and biological properties are typically inferior to those of the best green RSFPs. As a result, multi-color super-resolution microscopy based on red RSFPs remains characterized by a reduced performance [8,11–13].

At present, there is comparatively little mechanistic understanding of the photochromism in red RSFPs, making design of better-performing red RSFPs challenging. High-quality structural models of the different spectroscopic states could greatly enhance this understanding, though at present, structures are only available for rsTagRFP, asFP595 and IrisFP [14–16], and only rsTagRFP was fully characterized in both the fluorescent and non-fluorescent state [15]. The red-on asFP595 crystal structure could not be obtained directly due to its fast relaxation from on- to off-state, and point mutant variants had to be generated to determine the on-state structures [14]. The red-off IrisFP structures could not be clearly interpreted due to the presence of multiple chromophore conformations [16]. Based on the limited structural information and spectroscopic data of red RSFPs and analogy with green RSFPs, it has been proposed that on and off states adopt *cis* and *trans* configurations, respectively, and that the *cis-trans* isomerization is the molecular basis of photoswitching [17]. The spectroscopic data also proposes that the photoswitching is accompanied by protonation/deprotonation events at the chromophore tyrosine hydroxyl group [14,15,18], similar in concept to what has been determined for green RSFPs.

To expand on the photoswitching mechanism of red RSFPs, we studied the structural features underlying photoswitching in rsCherry (mCherry_E144V-I161S-V177F-K178W), one of the first reported red RSFPs [7]. However, we surprisingly found that solutions of purified rsCherry protein irreversibly lost their color at 4°C in the dark after two months. We identified molecular oxygen as the source of this fluorescence loss based on spectroscopic experiments. Through X-ray crystallography and mass spectrometry, we found that molecular oxygen reacts with the chromophore and leads to the cleavage of the chromophore *p*-hydroxyphenyl moiety in the process.

## 2. Materials and methods

### 2.1 Mutagenesis, protein expression and purification

Mutagenesis was performed using a modified QuikChange protocol [6]. All primers were designed by Quikchange Primer design tool (Agilent Technologies) and ordered from Integrated DNA Technologies (IDT). Used primers are listed in table S1. The plasmid (pRSETB) containing the protein of interest was transformed into *E. coli* JM109(DE3). A single colony was inoculated with 1L of Lysogeny broth (LB) medium supplemented with 100 μg/mL of ampicillin. The culture was shaken vigorously at 21°C (180-200 rpm) for 72 hours. The cells were then harvested and dissolved in ice-cold TN 100/300 buffer (100 mM Tris-HCl pH 7.4, 300 mM NaCl) before being lysed in a French Press. The resulting cellular debris was removed by centrifuging for 20 minutes at 8500 rpm, followed by incubating the supernatant with Ni-NTA agarose (Qiagen) for 40 minutes at 4°C. His-tag purification was performed in Pierce disposable polypropylene 5mL disposable columns (Thermo Fisher Scientific) with a wash solution containing TNi 100/300/20 (100 mM Tris-HCl pH 7.4, 300mM NaCl and 20 mM imidazole) and an elution solution containing TNi 100/300/250 (100 mM Tris-HCl pH 7.4, 300mM NaCl and 250 mM imidazole). Size exclusion chromatography was then performed in a HiLoad Superdex 200 pg 16/600 column coupled to an Äkta Prime (Cytiva) using a buffer consisting of TN 10/30 (10 mM Tris-HCl pH 7.4, 30mM NaCl). The eluted protein was then concentrated to the approximate concentration of 10 mg/ml using a Vivaspin 6 3,000 MWCO PES (Sartorius) membrane ultrafiltration tube.

### 2.2 Time-course spectroscopy in aerobic/anaerobic conditions

Anaerobic samples were prepared using a vacuum line. A process of sequential steps including freezing samples with liquid nitrogen, degassing by a vacuum line and thawing, was repeated three times until the pressure remained stable (approximately 10^-5^ bar). After that, absorption spectra and fluorescence of degassed and non-degassed samples were recorded using a Lambda 950 spectrophotometer (from Perkin Elmer) and FLS980 fluorometer (from Edinburgh), respectively.

### 2.3 Protein crystallization

The crystal of oxygen-degraded rsCherry was first grown at 21°C by the sitting drop vapor diffusion method in an aerobic environment at the Biomolecular Architecture laboratory, KU Leuven, Belgium. The mixture containing 0.75 μL of protein solutions (10 mg/mL) and 0.75 μL of precipitant (25% w/v PEG 8000, 0.1 M MES pH 6.5) was placed against a 70 μL reservoir. A colorless plate-like crystal of rsCherry was obtained after two months.

Anaerobic crystallization was performed at the Laboratory of Chemistry of Biological Processes, Collège de France, France. Microbatch and microseeding techniques [19] were used to grow crystals of rsCherry in a strictly oxygen-free environment. A serial dilution of the seed stock was prepared from microcrystals obtained in oxygen-degraded rsCherry. Protein solution (1 μL, 3-5 mg/mL) was mixed with 0.5 μL precipitant and 0.5 μL seed stock, then placed under a thin layer of Al’s oil. The crystallization cocktail was optimized with the best condition containing 18% w/v PEG 8000 and 0.1M MES pH 6, resulting in a bunch of plate-like colored crystals grown after a few days. Prior to flash cooling, the crystals were transferred to a similar condition as the reservoir with the addition of 25% glycerol as a cryoprotectant.

### 2.4 Data collection and refinement process

The X-ray diffraction data were collected at the beamlines X06DA (SLS, Switzerland) and Proxima2A (Soleil synchrotron, France). Crystals were diffracted under a nitrogen cryostream of 100 K using a wavelength of 1 Å. Diffraction images were indexed and integrated with XDS [20], and data reduction was performed using Aimless and Ctruncate (CCP4 7.0.078) [21]. The structures were solved by molecular replacement with Phaser (version 2.8.2) [22] in Phenix, using mCherry (PDB: 2H5Q [23]) as the search model. Before performing molecular replacement, the model of mCherry was edited by removing the chromophore as well as alternate conformers; and converting to isotropic temperature factors (PDBtools, Phenix). The structure was then refined and modeled by Phenix.refine (version 1.19.2_4158) [24] and Coot (version 0.9.6) [25], respectively. After the first refinement, all mutations along with the chromophore were modeled based on the electron density and difference maps. Dictionary files for the modified chromophores were created using JLigand [26] and manually adjusted using bond distances and angles restraints based on structures present in the Cambridge Structural Database (CSD) [27]. Waters were automatically added to the model using the “Update waters” option and were carefully double-checked (B-factors smaller than 80 Å^2^ and reasonably hydrogen bond distances ranging from 2.3-3.5 Å). Data collection and refinement statistics are listed in Table 1.

**Table 1.**
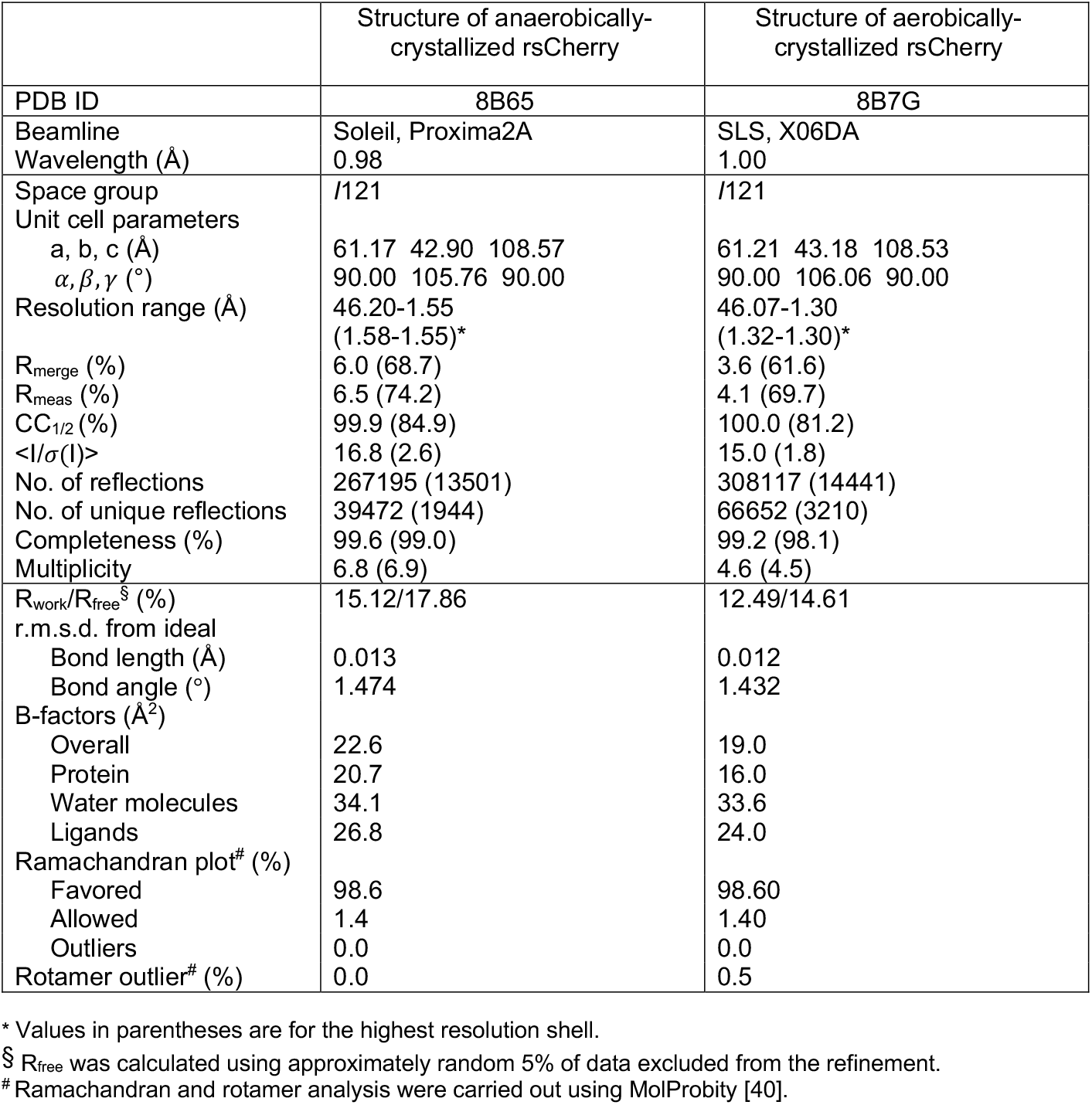
Data collection and Refinement statistics:

|  | Structure of anaerobically-crystallized rsCherry | Structure of aerobically-crystallized rsCherry |
| --- | --- | --- |
| PDB ID | 8B65 | 8B7G |
| Beamline | Soleil, Proxima2A | SLS, X06DA |
| Wavelength (Å) | 0.98 | 1.00 |
| Space group | I121 | I121 |
| Unit cell parameters |  |  |
| a, b, c (Å) | 61.17 42.90 108.57 | 61.21 43.18 108.53 |
| $\alpha, \beta, \gamma$ (°) | 90.00 105.76 90.00 | 90.00 106.06 90.00 |
| Resolution range (Å) | 46.20-1.55<br>(1.58-1.55)* | 46.07-1.30<br>(1.32-1.30)* |
| R<sub>merge</sub> (%) | 6.0 (68.7) | 3.6 (61.6) |
| R<sub>meas</sub> (%) | 6.5 (74.2) | 4.1 (69.7) |
| CC<sub>1/2</sub> (%) | 99.9 (84.9) | 100.0 (81.2) |
| <I/σ(I)> | 16.8 (2.6) | 15.0 (1.8) |
| No. of reflections | 267195 (13501) | 308117 (14441) |
| No. of unique reflections | 39472 (1944) | 66652 (3210) |
| Completeness (%) | 99.6 (99.0) | 99.2 (98.1) |
| Multiplicity | 6.8 (6.9) | 4.6 (4.5) |
| R<sub>work</sub>/R<sub>free</sub><sup>§</sup> (%) | 15.12/17.86 | 12.49/14.61 |
| r.m.s.d. from ideal |  |  |
| Bond length (Å) | 0.013 | 0.012 |
| Bond angle (°) | 1.474 | 1.432 |
| B-factors (Å<sup>2</sup>) |  |  |
| Overall | 22.6 | 19.0 |
| Protein | 20.7 | 16.0 |
| Water molecules | 34.1 | 33.6 |
| Ligands | 26.8 | 24.0 |
| Ramachandran plot<sup>#</sup> (%) |  |  |
| Favored | 98.6 | 98.60 |
| Allowed | 1.4 | 1.40 |
| Outliers | 0.0 | 0.0 |
| Rotamer outlier<sup>#</sup> (%) | 0.0 | 0.5 |

\* Values in parentheses are for the highest resolution shell.
<sup>§</sup> R<sub>free</sub> was calculated using approximately random 5% of data excluded from the refinement.
<sup>#</sup> Ramachandran and rotamer analysis were carried out using MolProbity [40].

The structures of rsCherry crystallized in anaerobic and aerobic conditions were deposited in the PDB with accession codes of 8B65 and 8B7G, respectively. Images were generated using the PyMOL Molecular Graphics System (version 2.5.2); Schrödinger, LLC.

### 2.5 Calculation of oxygen diffusion pathways by CAVER3

The potential tunnels in rsCherry and mCherry were calculated by using the CAVER 3.0 [28] as a PyMOL plugin. Before running calculations, the crystal structures of anaerobically-crystallized rsCherry (PDB: 8B65) and mCherry (PDB: 2H5Q) were prepared by removing all ligands and waters (except the chromophore); and only the alternative conformations with higher occupancies were retained. The analysis was performed for mCherry and rsCherry with the same default parameters (minimum probe radius: 0.9 Å; shell depth: 4 Å; shell radius: 3 Å and clustering threshold: 3.5). The chromophore **1** (PDB ligand ID: QYX and CH6 in rsCherry and mCherry respectively) was chosen as the starting point.

### 2.6 Native mass spectrometry

Freshly-purified and oxygen-degraded rsCherry was buffer exchanged with 50 mM ammonium acetate using spin columns and data was acquired using electrospray ionization (ESI) with a Q-TOF mass analyzer (Synapt G2 HDMS, Waters). Calibration of the system was performed using sodium iodide (2 mg/mL) with a mass range of 400-9000 m/z. The operating parameters for the spectral acquisition were as follows: capillary voltage, 1.8 kV; sample cone voltage, 60 V; extraction cone voltage, 2 V; source temperature, 37°C; desolvation temperature, 150°C; backing pressure, 5.26 mbar; source pressure, 2.08 e-3 mbar; Trap, 1.44 e-3 mbar. Spectra were acquired in the range of 900-8000 m/z in positive-ion V mode. The molecular mass of the recorded spectra was calculated using MassLynx software (MassLynx version 4.1) and deconvolution of the mass spectra was performed through MaxEnt 1. All the data were acquired in technical duplicates.

## 3. Results

### 3.1. rsCherry loses color in the presence of oxygen

We purified rsCherry after recombinant bacterial expression, obtaining bright red solutions. Full chromophore maturation was achieved after storing rsCherry for three additional days at 4°C in the dark (Fig. 1A, B). After that, the purified protein lost its color upon further storage under these conditions, with a clear decrease of both the absorbance at ~570 nm and the fluorescence at ~610 nm (Fig. 1A, B). The rsCherry absorbance and fluorescence loss was accelerated when incubating the sample at 37°C, following a single exponential decay rate (t_1/2 abs rsCherry_=30.42 h, t_1/2 fluo rsCherry_=28.33 h) (Fig. 1C, D). Based on our previous experience of mCherry fluorescence being quenched through chromophore modification by the reducing reagent β-mercaptoethanol (βME) [29], we reasoned that the fluorescence disappearance in rsCherry might have been caused by a compound present in the storage buffer that is able to modify the rsCherry chromophore. This compound turned out to be dissolved oxygen. While >90% of rsCherry fluorescence was lost after 6 days of incubation at 37°C, degassing the storage buffer prevented most of the fluorescence loss (Fig. 1C, D). Under the same conditions, the ancestor protein mCherry also showed an oxygen-dependent fluorescence decrease, but this process was markedly slower compared to rsCherry (t_1/2 fluo rsCherry_ =28.33 h, t_1/2 fluo mCherry_=83.94 h) (Fig. 1C, D).

**Fig 1.**
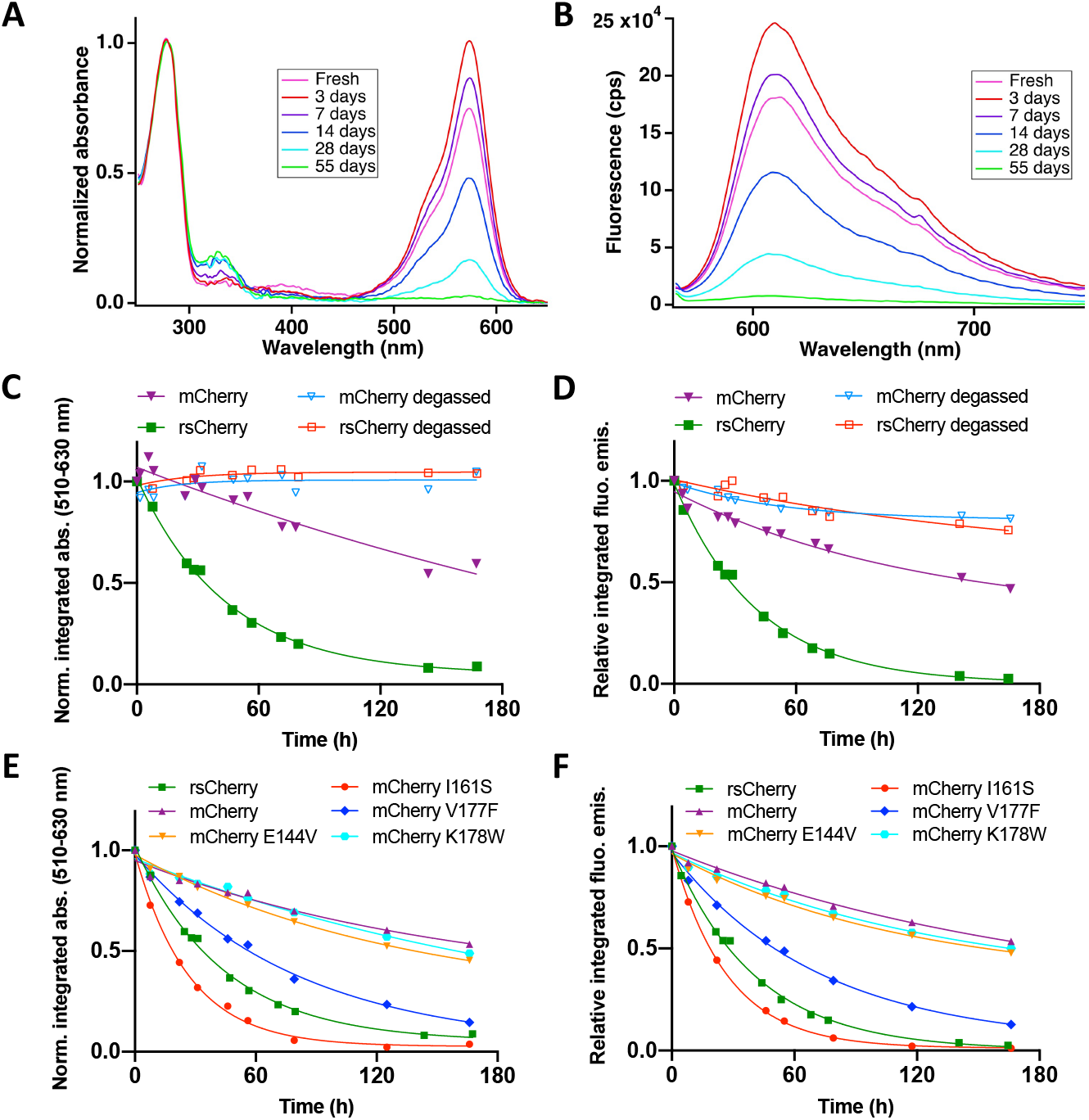
The loss of absorbance and fluorescence of rsCherry occurs spontaneously in aerobic conditions, and it is mainly caused by the I161S mutation. (A, B) Time-course of absorbance and fluorescence spectra of purified rsCherry at 4°C. (C, D) Time-course of normalized integrated absorption 510-630nm (C) and relative integrated fluorescence emission (D) of purified rsCherry and mCherry incubated at 37°C. The lines correspond to single exponential fittings of the absorbance/fluorescence decay. (E, F) Time-course of normalized integrated absorption 510-630nm (E) and relative integrated fluorescence emission (F) of purified mCherry and its single mutants mCherry_E144V/I161S/V177F/K178W incubated at 37°C. The lines correspond to single exponential fittings of the absorbance/fluorescence decay.

Given this difference in the fluorescence loss kinetics, it is likely that the mutations that were introduced while generating rsCherry from mCherry (E144V, I161S, V177F, K178W) lead to the enhanced oxygen-dependent loss of color. In mCherry single mutants, we observed that the I161S variant was even more oxygen sensitive than rsCherry (t_1/2 fluo mcherryI161S_=18.63 h) (Fig. 1E, F). Introducing the V177F mutation in mCherry also increased its susceptibility to oxygen-mediated degradation, but to a lesser extent than the I161S variant (t_1/2 fluo mCherryV177F_=49.83 h) (Fig. 1E, F). On the other hand, the E144V or K178W variants of mCherry, containing a single mutation located in the outer part of the protein, displayed no increase in the oxygen-induced degradation rate compared to mCherry (Fig. 1E, F). Conversely, we tested whether mutating S161I in rsCherry would cause a delay of its degradation by oxygen, but this effect was relatively small (t_1/2 fluo rsCherryS161I_=41.61 h) (Fig. S1), suggesting that the fast oxygen-mediated degradation in rsCherry requires the combination of two inner residues in the chromophore vicinity (I161S and V177F).

### 3.2. The observed degradation involves the loss of the *p*-hydroxyphenyl ring in the chromophore

To understand the molecular basis underlying the rsCherry fluorescence loss, we solved the X-ray structures of rsCherry crystallized under aerobic (PDB ID: 8B65) and anaerobic (PDB ID: 8B7G) conditions, at resolutions of 1.3 Å and 1.55 Å, respectively (Table 1). The anaerobically-crystallized rsCherry crystal displayed a noticeable red color, while the aerobically-crystallized rsCherry crystal was colorless (Fig. 2A, B). Both proteins crystallized in the same space group *I121* with almost identical unit cell parameters and one molecule in the asymmetric unit. In addition, the overall structures are very similar to that of mCherry (PDB: 2H5Q) [23] with root mean square deviation (r.m.s.d) values of 0.27 Å (aerobically-crystallized rsCherry) and 0.26 Å (anaerobically-crystallized rsCherry) when fitting 212 Cα atoms from residues 6 to 220.

**Fig 2.**
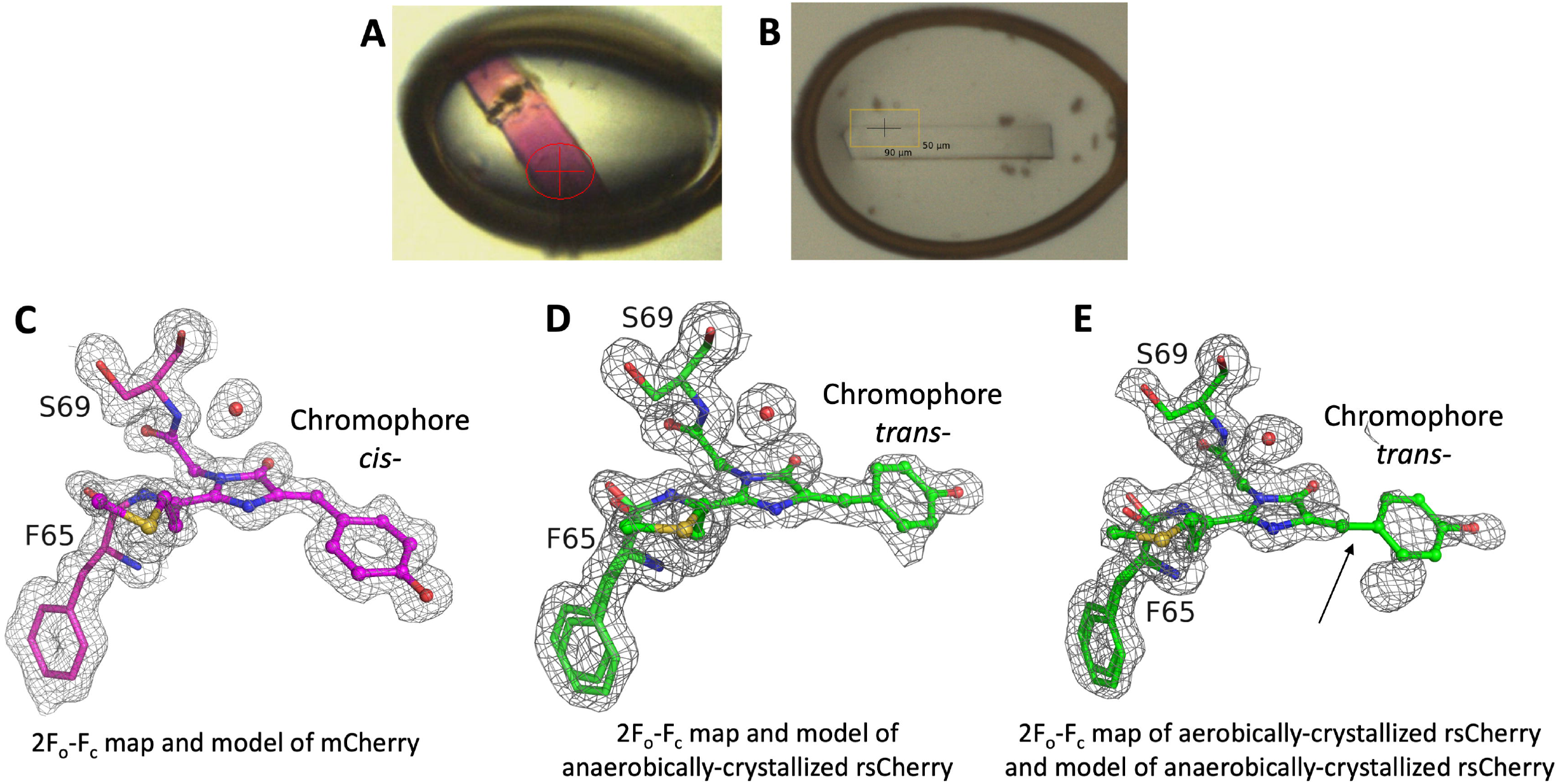
The crystal structures of rsCherry crystallized in anaerobic or aerobic conditions. Photographs of anaerobically-crystallized (A) and aerobically-crystallized (B) rsCherry crystals used for X-ray data collection, with their length being approximately 0.2 mm and 0.3 mm, respectively. (C-E) Comparison of 2Fo-Fc electron density maps contoured at 1 r.m.s.d of aerobically. and anaerobically-crystallized rsCherry structures. The chromophore **1** is depicted in ball-and-stick mode. (C) 2Fo-Fc map and model of the chromophore **1** (*cis* configuration) of mCherry (PDB ID: 2H5Q [23]). (D) 2Fo-Fc map and model of the chromophore **1** (*trans* configuration) of anaerobically-crystallized rsCherry (PDB ID: 8B65). (E) 2Fo-Fc map of the aerobically-crystallized rsCherry (PDB ID: 8B7G) superposed on the chromophore **1** (*trans* configuration) of anaerobically-crystallized rsCherry, showing the missing electron density for the *p*-hydroxyphenyl ring, indicated by an arrow.

We found that the chromophore of the anaerobically-crystallized rsCherry adopts a *trans* configuration, unlike the *cis* configuration that is observed in mCherry [7] (Fig. 2C, D). In aerobically-crystallized rsCherry, on the other hand, the chromophore lacks electron density where the *p*-hydroxyphenyl ring is expected to be (Fig. 2E), indicating that the *p-* hydroxyphenyl ring is cleaved during the oxygen-dependent process.

To confirm the oxygen-mediated loss of the chromophore *p*-hydroxyphenyl ring, we performed native mass spectrometry (MS) on freshly purified rsCherry and on rsCherry that had undergone oxygen-dependent degradation over three months. Freshly purified rsCherry displays a major peak at 30677 Da, which is the protein with the intact chromophore, and a second most abundant peak at 30697 Da which matches the expected mass for rsCherry with incomplete chromophore maturation (Fig. 3A, S2). In the case of aerobically-degraded rsCherry, two peaks of similar intensity are found: one at 30587 Da, also present as a minor peak in freshly purified rsCherry, and a unique one at 30605 Da (Fig. 3A, S2).

**Fig. 3.**
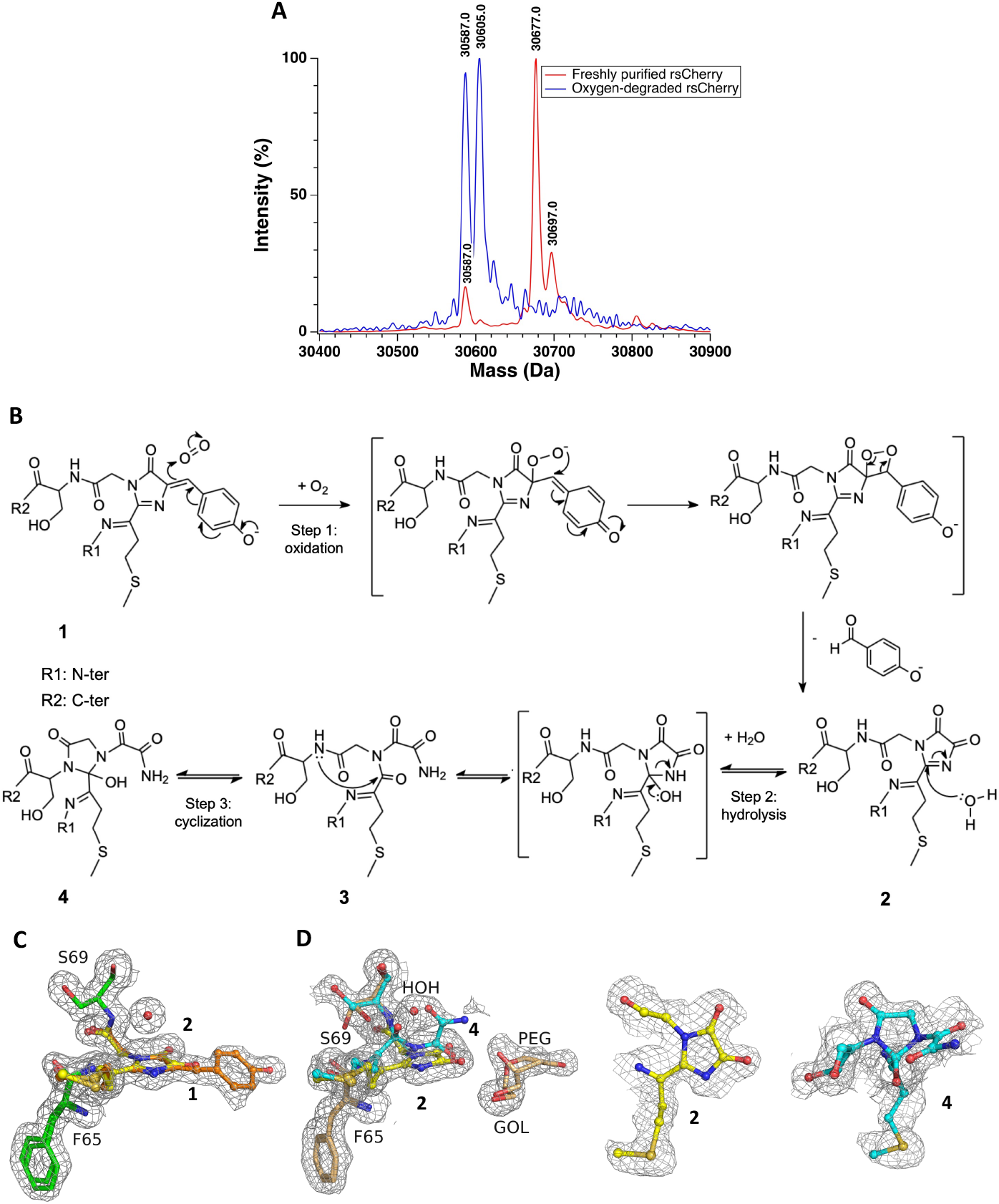
(A) Deconvoluted mass spectra of the freshly purified and oxygen-degraded rsCherry obtained by native mass spectrometry. (B) Proposed mechanism for the multi-step oxygen-mediated degradation in rsCherry. Freshly purified rsCherry chromophore structure **1** (Mw = 30677 Da) is first oxidized by oxygen dissolved in the buffer, gradually forming **2**, leading to a mass loss of 90 Da which corresponds to the missing *p*-hydroxyphenyl ring (the degraded species Mw = 30587 Da). Due to its high reactivity, **2** is assumed to be further hydrolyzed to form **3**. Compound **3** possibly undergoes an intramolecular nucleophilic attack by the nitrogen atom of the adjacent residue S69, resulting in **4**. The structure **4** is expected to have a mass of 30605 Da, matching with the peak at 30605 Da in the spectrum. (C, D) The models and electron density maps of the anaerobically-crystallized (PDB: 8B65) and aerobically-crystallized rsCherry (PDB: 8B7G), respectively. Chromophore structures **1**, **2**, **4** are represented in ball-and-sticks; and colored orange, yellow and cyan, respectively. Protein backbones of the anaerobically and aerobically-crystallized rsCherry are colored green and light orange respectively (C) The final model and 2Fo-Fc maps contoured at 1 r.m.s.d of anaerobically-crystallized rsCherry. The occupancies of **1** and **2** at the chromophore position converged towards 0.33 and 0.67, respectively. (D) The final model and 2Fo-Fc maps contoured at 0.5 r.m.s.d. of all (left) and individual (middle and right) structures of **2** and **4** in aerobically-crystallized rsCherry. Structure refinement with structures **2** and **4** led to occupancies of 0.68 and 0.32, respectively. PDB ligand IDs of **1**, **2** and **4** in the structures of rsCherry are QYX, Q2K and QIP respectively.

The combination of the X-ray crystallography and native MS data suggests the following mechanism of rsCherry chromophore destruction by oxygen: upon exposure to air, oxidative cleavage of the methylene bond linking the 4-imidazolinone to the *p*-hydroxyphenyl ring results in an imidazole-4,5-dione ring (Fig. 3B, structures **1**, **2**). This degradation product has a 90 Da mass decrease compared to the original chromophore, matching with the 30587 Da peak observed in both freshly purified and oxygen-degraded rsCherry (Fig. 3A). We interpret the presence of the minor 30587 Da peak in the freshly purified sample as arising through degradation that already occurred during protein expression and/or purification. We included structures **1** and **2** into our X-ray analysis of the anaerobically-crystallized rsCherry model, resulting in a model with occupancies of 0.67 and 0.33 for structures **1** and **2**, respectively (PDB ID: 8B65, Fig. 3C). We then tried to model the structure of aerobically-crystallized rsCherry with only structure **2**, resulting in a good fit to the observed electron density, though this resulted in an incomplete occupancy (0.78) of this structure. This indicated that additional structures are present in the chromophore region (Fig. S3).

One of the remaining structures is related to the 30605 Da peak of oxygen-degraded rsCherry, which has an 18 Da mass increase as compared to the protein with chromophore structure **2**. Looking at the electron density map in chromophore region, there is an unmodelled density with a cyclic shape at S69 which suggests that another five-membered ring is present in the degraded chromophore (structure **4,** Fig. 3B,D, S3B). Therefore, we propose that, upon formation of structure **2**, a nucleophilic addition of a water molecule to the C=N bond of the imidazole-4,5-dione of structure **2** results in a ring opening to form structure **3** [30,31] (Fig. 3B). The presence of electron-withdrawing imine groups in **3** can make the newly formed carbonyl carbon atom more partially positive and susceptible to an intramolecular nucleophilic attack by the amide nitrogen atom of S69, resulting in the formation of structure **4** (Fig. 3B). However, structure refinement with chromophore structures **2**, **3** and **4** leads to an occupancy of only 0.14 for structure **3**, suggesting that it quickly reacts to form **4** (Fig. S4). Therefore, we finally refined the model of aerobically-crystallized rsCherry with structures **2** and **4**, obtaining occupancies of 0.68 and 0.32, respectively (PDB ID: 8B7G). Despite improvement in the map, the resulting F_o_-F_c_ map still contains some residual difference density, suggesting further or alternative modifications in the chromophore area which cannot be interpreted clearly in the X-ray structure (Fig. S5, S6).

### 3.3. Additional oxygen accessibility to the chromophore could explain the differences in degradation between mCherry and rsCherry

rsCherry (mCherry_E144V-I161S-V177F-K178W) and mCherry are closely related yet show pronounced differences in their resistance to oxygen-induced degradation. We hypothesized that this contrast likely arises from the different accessibility of the chromophore to oxygen. To examine this more systematically, we used CAVER 3.0 [28] to predict potential transport tunnels for small molecules in each protein (Fig. 4A, B). This analysis found that mCherry and anaerobically-crystallized rsCherry structures have two similar tunnels, but rsCherry also has two additional tunnels (Fig. 4A, B; Table S2, S3). The presence of the extra tunnels in rsCherry could be explained by the structural differences that are induced by the I161S and V177F mutations. The I161S mutation favors the presence of the *trans*-chromophore in rsCherry by hydrogen bonding between S161 and the chromophore hydroxyl groups. The V177F mutation introduces a bulky residue in the chromophore pocket that forces the movement of several residues to avoid steric clashes. Combined, these two mutations lead to several conformational changes of residue side chains in the chromophore pocket, such as S146, E148, Q163 and K70, resulting in an altered hydrogen bond network (Fig. 4C, D). In particular, the movement of K70 side chain is crucial regarding oxygen accessibility towards its attack at the chromophore methylene bridge. Due to the presence of the *trans*-chromophore observed in rsCherry, K70 moves approximately 3.0 Å away from the imidazolinone ring in comparison to the same residue in mCherry (Fig. 4E). This creates an extra space where an oxygen molecule could reside, increasing the chances of oxygen-mediated chromophore degradation at the chromophore methylene bridge. On the other hand, the chromophore of mCherry is in the *cis* configuration and K70 is closer to the chromophore, which protects the chromophore from direct oxygen accessibility (Fig. 4E). A similar K70 side chain displacement has only been observed upon the thio-adduct formation between the mCherry chromophore methylene bridge and β-mercaptoethanol (βME) after this protein was treated with βME [29].

**Fig. 4.**
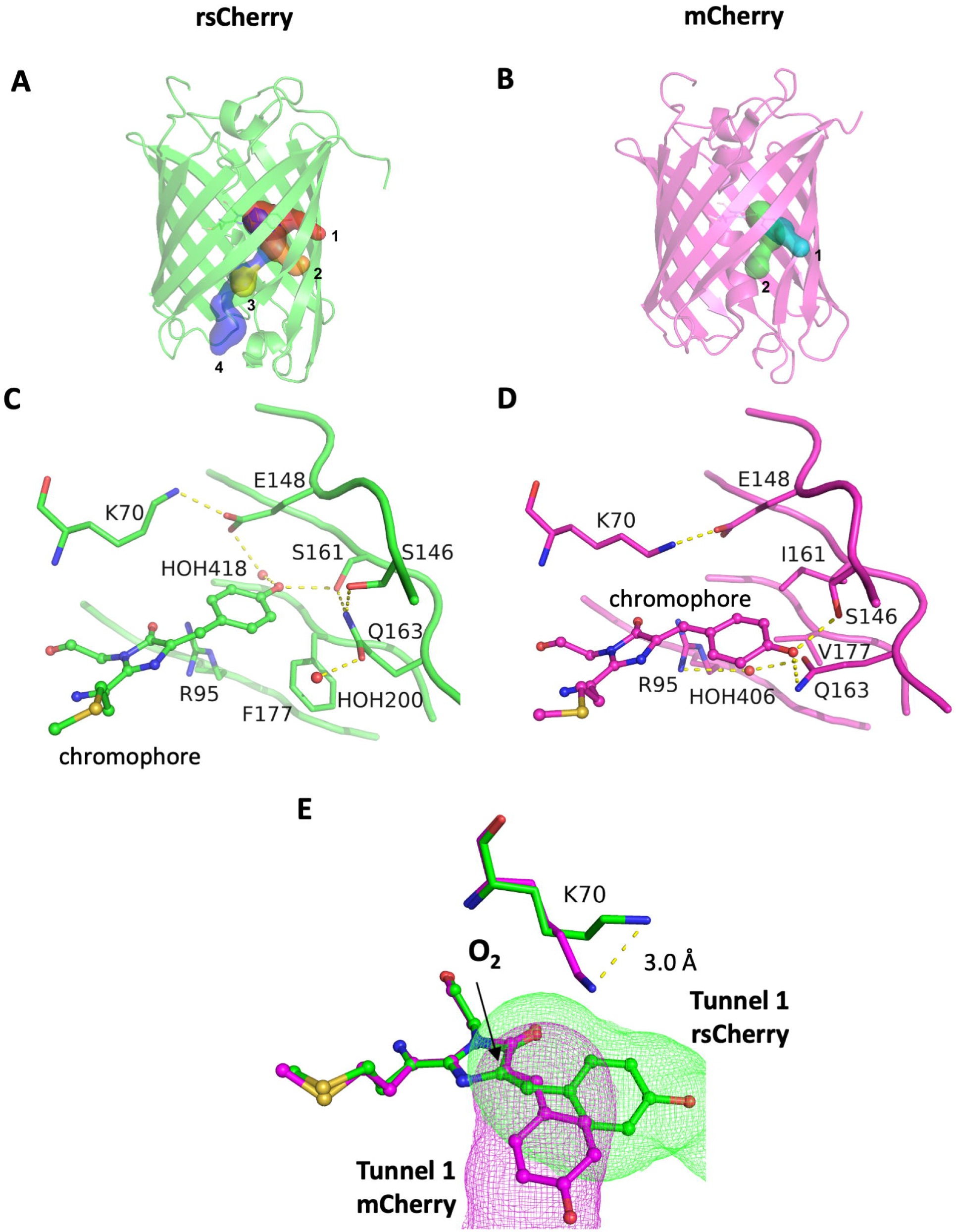
Comparison of oxygen accessibility and structures of anaerobically-crystallized rsCherry (green) and mCherry (magenta). (A,B) The structures of rsCherry (A) and mCherry (B) are shown as cartoons and tunnels predicted by CAVER 3.0 are visualized as surface. The tunnels are numbered according to their priority estimated by CAVER 3.0. Compared to mCherry, two additional tunnels (1 and 2) are identified in rsCherry. (C, D) Hydrogen bond network in the chromophore pocket of mCherry and rsCherry. The combination of I161S and V177F mutations in rsCherry causes movements of E148, S146, Q163 and K70 side chains, resulting in chromophore microenvironment differences between mCherry and rsCherry. (E) The crucial role of K70 in oxygen accessibility for degradation. In rsCherry, K70 displays a shift of 3.0 Å compared to mCherry, creating sufficient space for oxygen attack at the methylene bridge of the chromophore. The conformation of K70 in mCherry does not support the presence of similar tunnels as in rsCherry. The chromophores are depicted in ball-and-stick and tunnels are represented as isomesh. Tunnels of mCherry and rsCherry are colored pink and green, respectively. The arrow indicates the position where oxygen attacks the rsCherry chromophore.

Remarkably, we observed the presence of a glycerol molecule (PDB ligand ID: GOL) close to the chromophore in the aerobically-crystallized rsCherry structure with an occupancy of 0.36 (Fig. 3D). This glycerol originates from the cryoprotectant solution that only contacted the rsCherry crystal very quickly (less than a minute) before flash freezing, confirming transport pathways for small molecules to access the rsCherry chromophore and release reaction products to the surrounding buffer solution (Fig. S7).

## 4. Discussion

Oxygen is a required molecule for the chromophore maturation in fluorescent proteins. In the case of rsCherry, however, we observed that oxygen causes the degradation of the mature chromophore, leading to a loss of absorption and fluorescence. While the structural features in rsCherry chromophore region make it particularly prone to oxygen-mediated degradation, oxygen also caused mCherry fluorescence loss over time, albeit to a lesser extent. This result suggests that other FPs could be subject to this oxygen-mediated fluorescence decrease, with one key parameter being the accessibility of the methylene bridge to molecular oxygen.

Compared to other modifications in other fluorescent protein chromophores [31–34], three aspects make this chemical modification in rsCherry special:

*First*, the degradation in rsCherry is an oxygen-mediated process that is light-independent. Our study shows that the freshly purified rsCherry loses its absorbance and fluorescence spontaneously in the dark, and that removing oxygen from the buffer greatly delays its degradation. The involvement of molecular oxygen in FP bleaching has been previously observed in IrisFP at low light intensity [35]. However, the degradation in rsCherry is likely to proceed differently from the low-intensity photobleaching in IrisFP, as the IrisFP chromophore was intact and showed no modifications [35]. This type of bleaching in IrisFP was presumably caused by the sulfur oxidation of M159 and C171 side chains, raising the pKa of the chromophore and trapping the IrisFP chromophore in the dark protonated state [35].

*Second*, the initial cleavage position at the bond Cα-Cβ leading to the removal of the *p-* hydroxyphenyl ring in the chromophore, is very uncommon. Most chromophore modifications in GFP or RFP homologs happen at the imidazolinone ring and/or at the acylimine link of the red chromophore [31,32,36]. There are also a few examples of chemical reactions occurring at the bond Cα-Cβ bridging two cyclic moieties of the chromophore but via completely different pathways. For instance, the Cα-Cβ bond in the phenol side chain of mTagBFP can be oxidized from a single bond to a double bond, generating TagRFP [37], but the *p*-hydroxyphenyl ring cleavage does not occur. Another example of chemical modification at the same location is the reaction of mCherry and βME through a Michael addition reaction that forms a thio-adduct at the Cβ atom of the mCherry chromophore [29], leading to a loss of absorbance and fluorescence. However, in contrast to the oxidation of rsCherry by molecular oxygen, the reaction of mCherry with βME is reversible, without *p*-hydroxyphenyl ring cleavage, and there is an additional absorbance peak arising at 410 nm. Compared to the chromophores of mTagBFP and mCherry-βME, the resulting products after chromophore degradation in rsCherry, structures **2** and **4**, do not absorb at ~410 nm, and only a ~320nm absorbance peak is observed. Structure **4** has no longer a conjugation system and thus is not supposed to absorb light at 320 nm. In structure **2**, which still retains a conjugation system and thus can presumably absorb UV light, the loss of the *p*-hydroxyphenyl moiety could explain the shorter wavelength absorption maxima compared to the chromophores of TagBFP or mCherry-βME, where the *p*-hydroxyphenyl moiety is still present. An absorbance shift to shorter wavelengths has also been observed in a special GFP variant with an engineered ASG chromophore forming tripeptide. This nonfluorescent variant absorbs at ~385nm, but upon heat treatment, the protein backbone breaks at the bond between the imidazolinone ring of the chromophore and the N-terminal leading amino acid residue, resulting in an absorbance shift to ~340nm [32].

*Third*, our obtained results demonstrated that the degradation of the rsCherry chromophore is not a side reaction of the maturation process but occurs after the full formation of the red chromophore, strikingly with only a cofactor - oxygen - that is indispensable for chromophore maturation. This post-maturation modification is distinct from the other reported modified structures of FPs chromophores. While some red chromophores of RFPs also undergo chemical modifications after maturation, such as a backbone cleavage at the acylimine bond in DsRed [38] or chromophore damage in KillerRed [39] and IrisFP [35], these modifications only occur upon heating or upon irradiation with green or cyan light.

## 5. Conclusion

This study demonstrates a new chemical reactivity in fluorescent proteins, where molecular oxygen - a cofactor needed for FP chromophore maturation - is the causative agent that degrades the chromophore of rsCherry (mCherry_E144V-I161S-V177F-K178W). The two inner mutations (I161S and V177F) lead to a series of structural rearrangements that increase the rate of oxygen-induced chromophore degradation in rsCherry. Among these mutations, the I161S mutation plays the most crucial role in promoting the oxygen-dependent degradation, as it probably induces the chromophore *trans* configuration and the K70-related displacement which allows oxygen to easily access the chromophore moiety and degrade it. While the structural features in the rsCherry chromophore region make it particularly prone to oxygen-mediated degradation, oxygen also causes mCherry fluorescence loss over time, albeit to a lesser extent. These results raise the possibility of other FPs being equally subject to this oxygen-mediated fluorescence loss once purified and stored under oxygen-containing conditions.

## 6. Acknowledgments

We thank Wim Dehaen, Department of Chemistry, KU Leuven, for insightful discussion. T.Y.H.B. and L.V.M. thank the staff of the beamline X06DA at the Swiss Light Source (Switzerland) and Proxima2A at Soleil synchrotron (France) for their assistance with the data collection. T.Y.H.B. acknowledges support by grants from the Vietnamese government (Project 911) and Schlumberger Foundation (Faculty for the Future program). This project has received funding from the European Union’s Horizon 2020 research and innovation programme under the Marie Skłodowska-Curie grant agreement No 101030525 (to B.P.). This work was supported by the FWO via grant G090819N, the European Research Council via grant 714688 NanoCellActivity, and the KU Leuven via grant C14/17/111.

## 7. Author contributions

**T.Y.H.B.**: Conceptualization, Data curation, Formal analysis, Investigation, Methodology, Validation, Visualization, Writing – original draft; Writing – review & editing, Funding acquisition

**E.D.Z.**: Conceptualization, Data curation, Formal analysis, Validation, Visualization, Writing – review & editing.

**B.M.**: Conceptualization, Formal analysis, Methodology, Validation, Visualization, Writing – review & editing.

**L.P.**: Investigation, Writing – review & editing

**B.Y.S.**: Investigation, Formal analysis, Writing – review & editing

**A. E.**: Resources, Supervision, Writing – review & editing

**M.F.**: Resources, Supervision, Writing – review & editing

**L.V.M**.: Conceptualization, Methodology, Validation, Resources, Supervision, Project administration, Writing – review & editing

**P.D**.: Conceptualization, Methodology, Validation, Resources, Supervision, Project administration, Writing – original draft, Writing – review & editing, Funding acquisition

**B. P**.: Conceptualization, Formal analysis, Methodology, Validation, Supervision, Writing – original draft, Writing – review & editing, Funding acquisition

