## Supplementary information for "Oxygen-induced chromophore degradation in the photoswitchable red fluorescent protein rsCherry"

### Supplementary Figures

#### Supplementary figure 1: Effects of mutation S161I on oxygen-mediated degradation in rsCherry

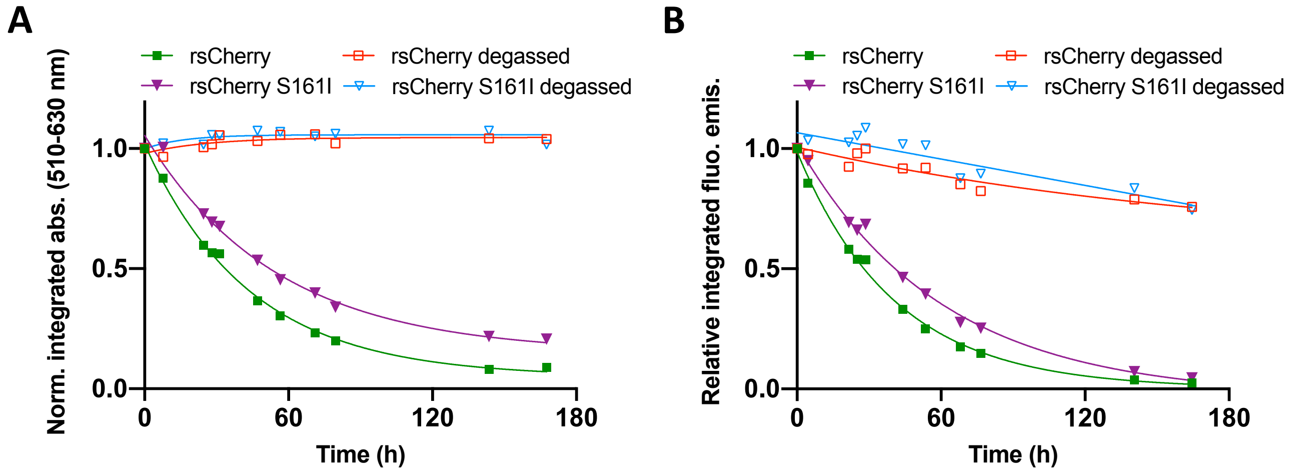


**Fig. S1.** Normalized integrated absorption (510-630 nm) and relative integrated fluorescence emission as a function of time of rsCherry and rsCherry S161I, incubated at 37°C. The lines correspond to single exponential fittings of the absorbance/fluorescence decay.

### Supplementary figure 2: Native mass spectrometry

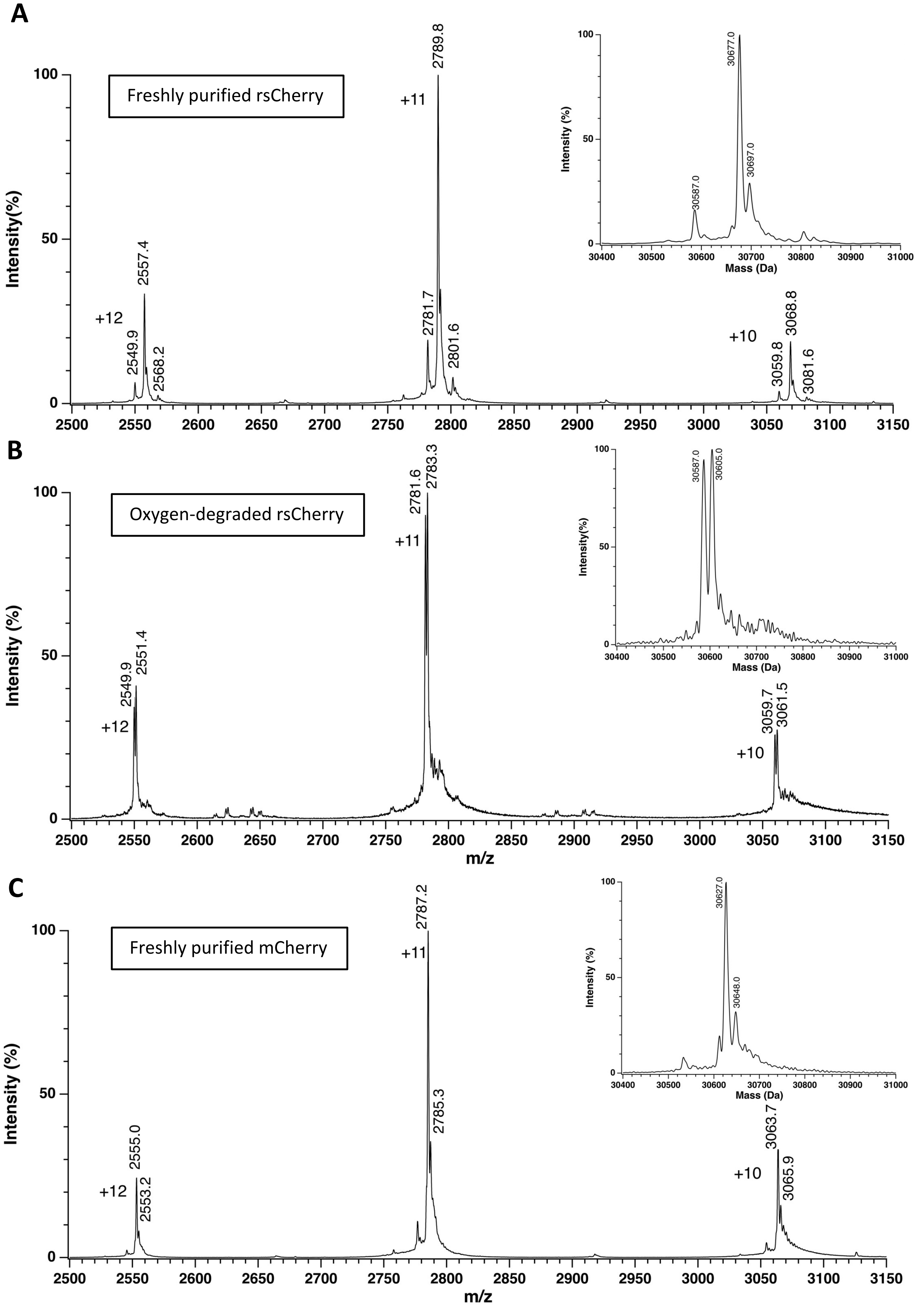

**Fig. S2.** Native mass spectra acquired on an ESI-mass spectrometer with Q-ToF mass analyzer (Synapt G2, Waters). Charge states are shown. The insets represent the deconvoluted mass in Da of (A) freshly-purified rsCherry [15 µM], (B) oxygen-degraded rsCherry [15 µM], and (C) freshly-purified mCherry [15 µM]. Note that the observed masses of both mCherry and freshly-purified rsCherry were approximately 15 Da higher than their theoretical masses due to modifications that occurred during native-MS experiments of intact proteins which are not caused by the studied degradation. The mass difference between mCherry and the freshly-purified rsCherry (functional protein) is 50 Da, in agreement to the four amino acid residues that are different in rsCherry compared to mCherry.

**Supplementary figure 3: The structure refinement of aerobically-crystallized rsCherry with the chromophore structure 2**

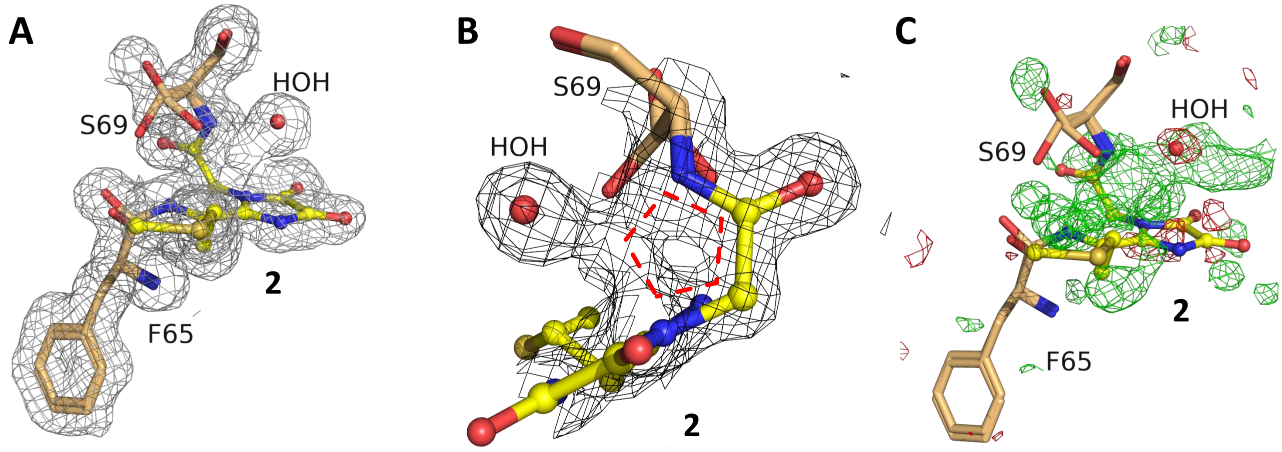


**Fig. S3:** Electron density of the aerobically-crystallized rsCherry model that was refined with only the chromophore structures **2** (occupancy 0.78). (A) $2F_o$-$F_c$ map contoured at 0.5 r.m.s.d at the chromophore position. (B) Closer view of electron density surrounding the chromophore **2** and S69 in the $2F_o$-$F_c$ map contoured at 1 r.m.s.d. The residual electron density clearly shows the shape of an extra five-membered ring (indicated as red dashed lines). (C) The $F_o$-$F_c$ map contoured at 3 r.m.s.d, showing many positive residual peaks, suggesting the presence of additional structures at the chromophore position.

**Supplementary figure 4: The structure refinement of aerobically-crystallized rsCherry with three chromophore structures 2, 3, 4.**

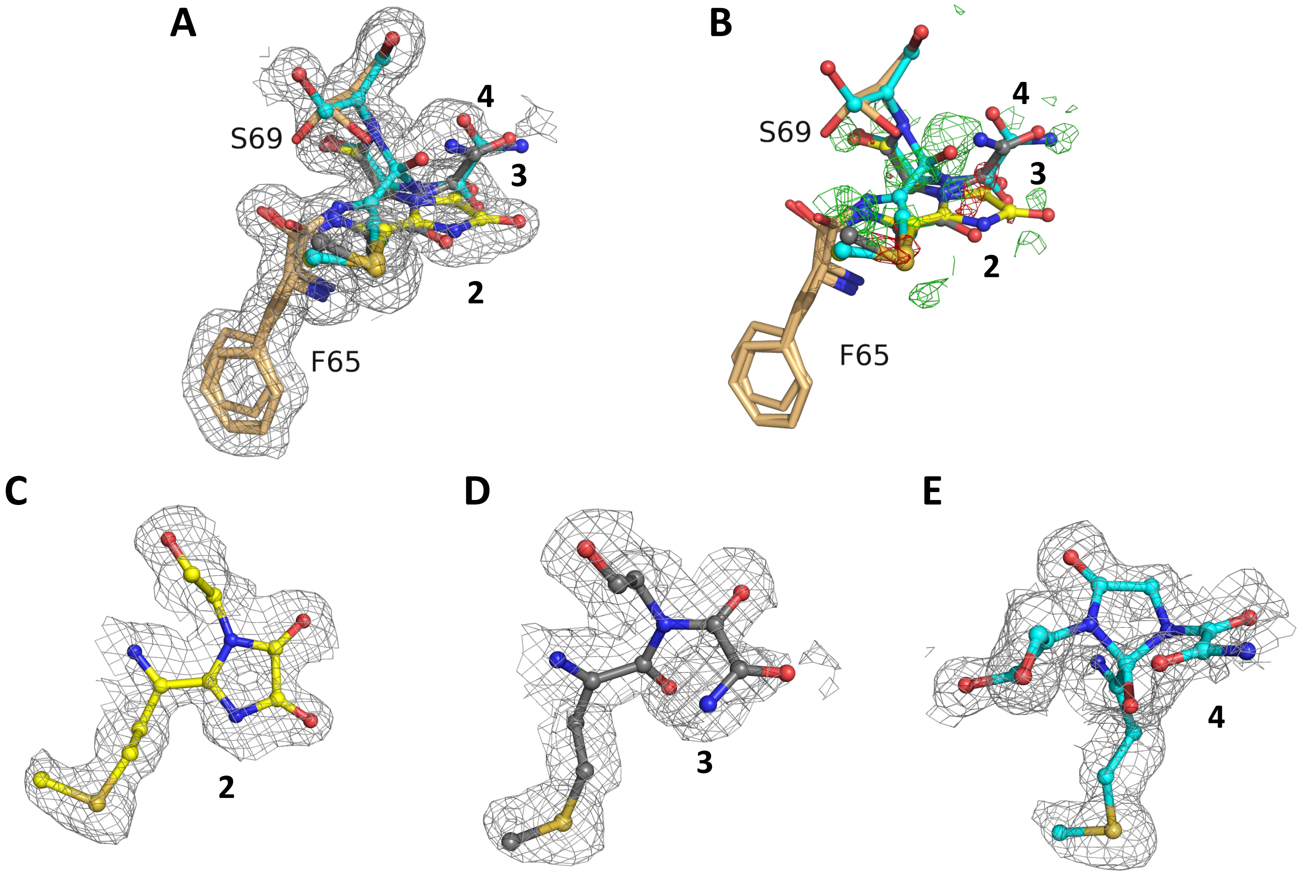


**Fig. S4.** Electron density maps calculated with the aerobically-crystallized rsCherry model refined with three chromophore structures **2**, **3**, **4** that were formed via the most probable chromophore modifications. The oxygen-degraded rsCherry was modeled with the combination of **2**, **3** and **4** with their occupancies of 0.54, 0.14 and 0.32 respectively. The maps do not show much improvement compared with the structure of aerobically-crystallized rsCherry modeled by **2** and **4** only (Fig. 3 and Fig. S6C,D). The low occupancy of **3** also indicated its ambiguous presence in the X-ray structure. (A, B) The complete model with the three chromophore types superposed on the $2F_o$-$F_c$ and $F_o$-$F_c$ map. (C, D, E) The $2F_o$-$F_c$ map and the individual chromophore structures **2**, **3**, **4**. $2F_o$-$F_c$ maps (A, C, D, E) are contoured at 0.5 r.m.s.d and the $F_o$-$F_c$ map (B) is contoured at 3 r.m.s.d.

**Supplementary figure 5: Proposed pathways for rsCherry chromophore degradation**

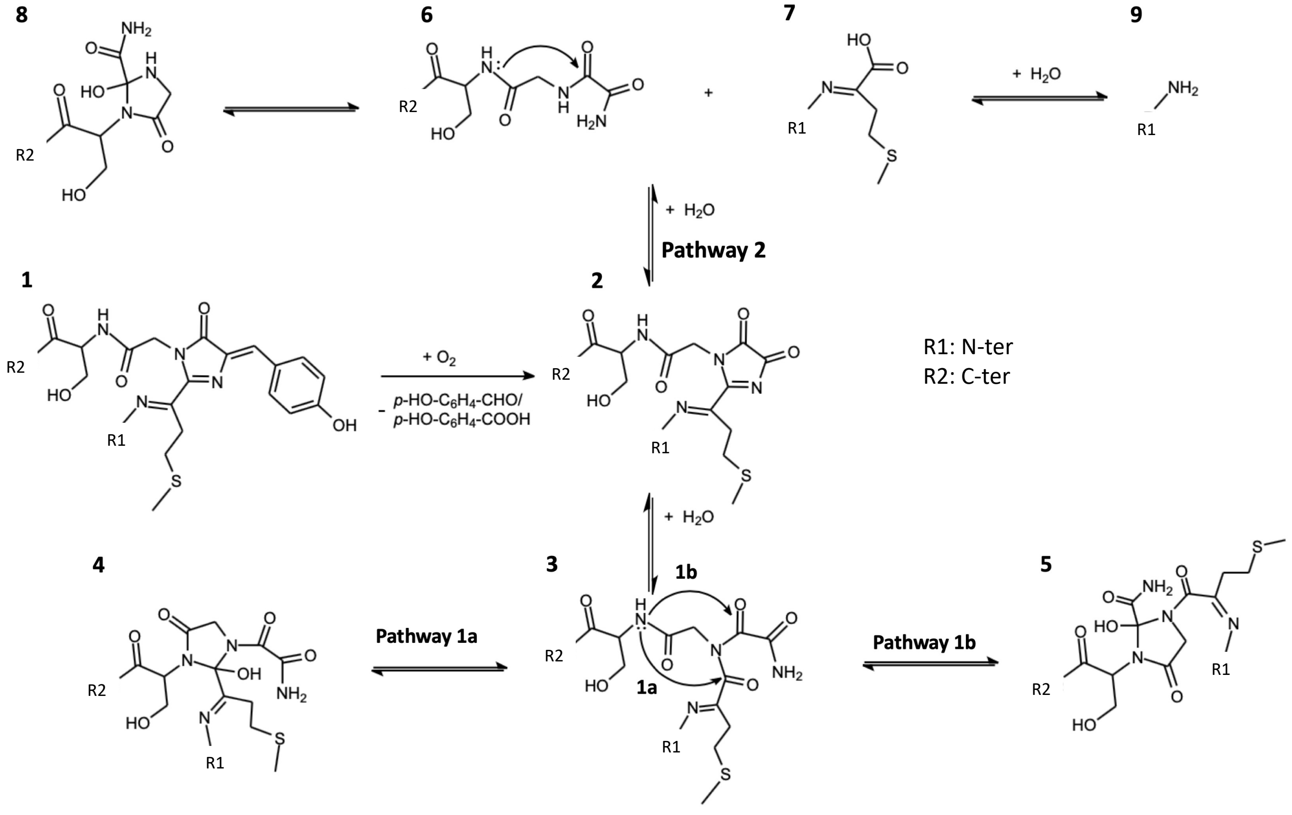


**Fig. S5.** Possible pathways for the degradation of purified rsCherry. We hypothesize that chromophore degradation starts oxidation of the chromophore **1**, leading to the cleavage of *p*-hydroxyphenyl and the formation of **2.** Subsequent hydrolysis of the imidazole-4,5-dione ring results in **3**, followed by an intramolecular nucleophilic attack of the N atom in S69 on one of the two reactive carbonyl groups in **3** to form **4** or **5** via pathway 1a or 1b, respectively. In another pathway (pathway 2), backbone hydrolysis of **2** results in the formation of **6** and **7**, which could further convert into **8** and **9** (based on the degradation observed in Blue102 [1]). Based on the mass spectroscopy and crystallographic analysis, pathway 1a is suggested as the most probable.

**Supplementary figure 6: Possible models for the aerobically-crystallized rsCherry structures**

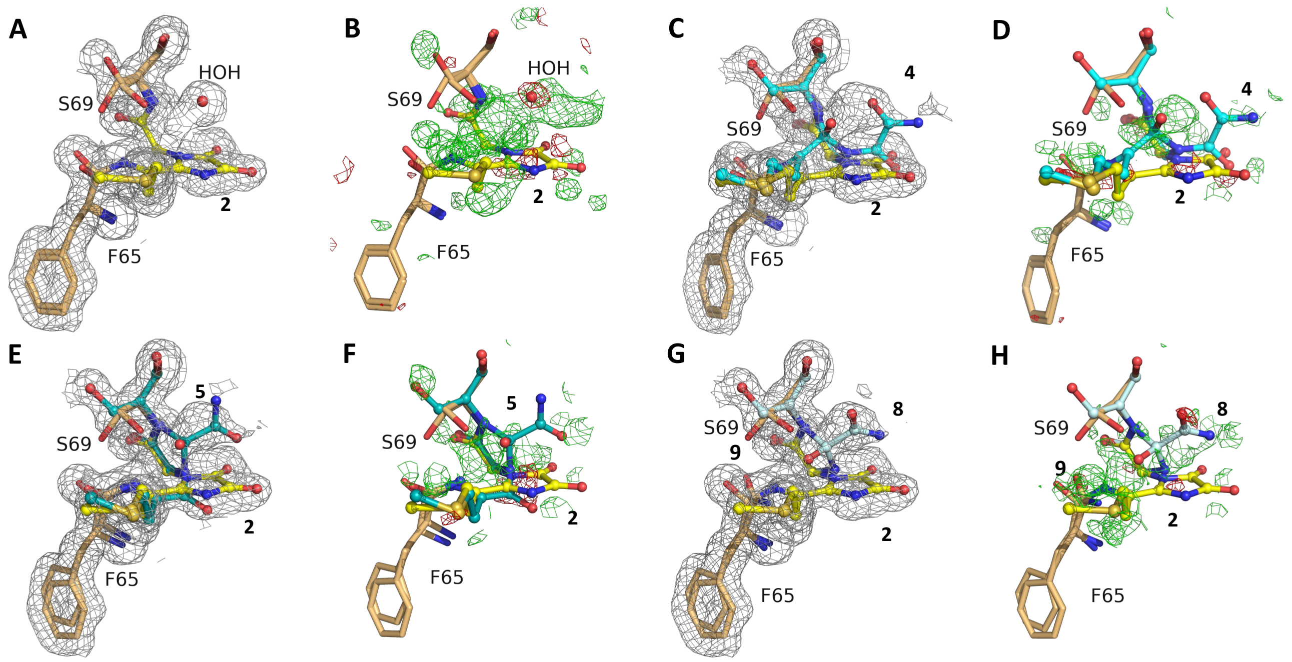


**Fig. S6.** Electron densities calculated using different modified structures of the chromophores in the aerobically-crystallized rsCherry structure. (A, C, E, G) $2F_o$-$F_c$ maps are contoured at 0.5 r.m.s.d; (B, D, F, H) $F_o$-$F_c$ maps are contoured at 3 r.m.s.d. (A, B) Electron density calculated with only **2** at an occupancy of 0.78 modeled at the chromophore position. (C, D, E, F) Electron density calculated with a combination of **2** and **4** or **2** and **5**, respectively. Degradation pathways 1a and 1b (Fig. S4) would result in the structure **2**, associated with a mass loss of 90 Da, and other species **3**, **4** or **5** that have similar mass losses of 72 Da compared to the functional protein. These degradation pathways agree better with the native mass spectrometry than alternative pathway 2. Although the refinement with **2**, **4** gave similar R-factors as the combination of **2**, **5** (see Table S4), the electron density appeared better for **2, 4** and we observe a higher agreement with the geometry restraints. Therefore, pathway 1a was assumed to mainly occur and we thus modelled **2**, **4** in the structures of aerobically-crystallized rsCherry. However, the alternative pathways cannot be excluded. (G, H) Electron density calculated using **2** and degradation products for pathway 2. Similar reactions were previously reported for Blue102 and Fast-FT [1]. As a result of degradation, a partial backbone cleavage occurs between residue F65 and the chromophore leading to the formation of **8** and **9**. After refinement, the occupancies of **2** and **8** converged to 0.66 and 0.34 respectively, which means that approximately 34% of the protein was cleaved to form **8** and phenylalanyl amide **9**. However, little backbone cleavage was observed in oxygen-degraded rsCherry when loaded on SDS-PAGE (data not shown), indicating that backbone cleavage-based pathway 2 is not prominent.

**Supplementary figure 7: The presence of a glycerol molecule in the chromophore pocket in the aerobically-crystallized rsCherry structure.**

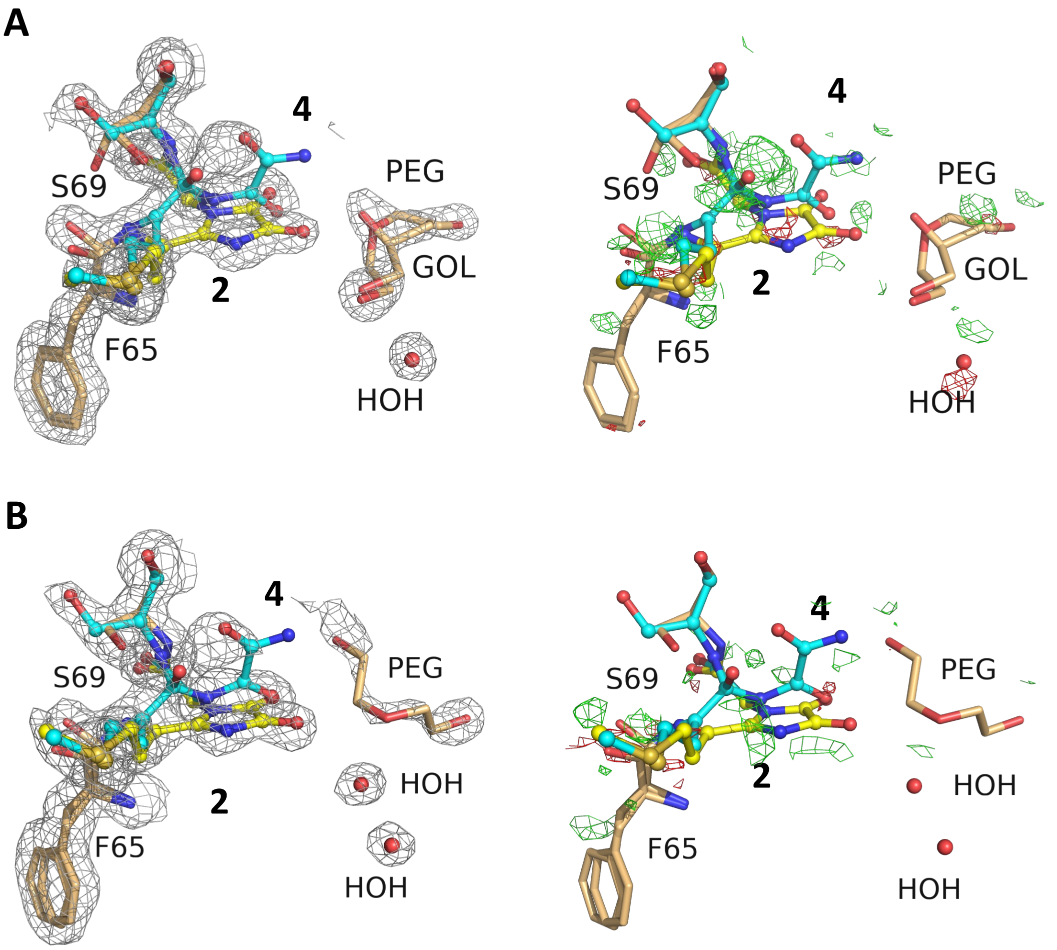


**Fig. S7.** Comparison between the electron density maps of two aerobically-crystallized rsCherry crystals with glycerol (A) and without glycerol (B) in the cryoprotectant solution. The two crystals were grown in the same conditions. (A) Electron density maps of the crystal soaked in a cryoprotectant solution containing 25% glycerol (resolution of 1.3 Å). (B) Electron density maps of the crystal soaked in a cryoprotectant solution containing 20% PEG 400 (resolution of 1.55 Å). The differences in the electron density maps at the chromophore pocket between the two structures indicate the effect of glycerol in the cryoprotectant solution. In structure **A**, one molecule of glycerol and one molecule of diethylene glycol appear close to the chromophore with occupancies of 0.33 and 0.42, respectively; while structure **B** only shows a diethylene glycol molecule (occupancy of 0.50). Diethylene glycol was formed during crystallization due to polyethylene glycol degradation in the mother liquor, while glycerol can only originate from the cryoprotectant solution. (Left) $2F_o$-$F_c$ maps contoured at 1 r.m.s.d; (Right) $F_o$-$F_c$ maps contoured at 3 r.m.s.d.

### Supplementary tables

### Table S1. List of used primers

| Name of primer | Primer sequence |
|---|---|
| **mCherry_E144V_fw** | CATGGGCTGGGTGGCCTCCTCCG |
| **mCherry_I161S_fw** | CCTGAAGGGCGAGGGCAAGCAGAGGCT |
| **mCherry_V177F_fw** | CCACTACGACGCTGAGTTCAAGACCACCTCCAA |
| **mCherry_K178W_fw** | CTACGACGCTGAGGTCTGGACCACCTACAAGGCC |
| **mCherry_V177F,K178W_fw** | CCACTACGACGCTGAGTTCTGGACCACCTACAAGGCC |

### Table S2. Characteristics of tunnels identified in mCherry by using CAVER 3.0

| **Tunnel ID** | **Average Bottleneck radius (Å)** | **Priority** | **Bottleneck residues** | **Residues in distance ≤ 3.0 Å from the bottleneck (sorted from the closest)** |
|---|---|---|---|---|
| 1 | 1.138 | 0.57319 | W143 | A145, K198, E144, L199, D200, K166 |
| 2 | 0.986 | 0.54044 | L199 | W143, S62, D200, E144, K198, Q163, P63, G142 |

### Table S3. Characteristics of tunnels identified in rsCherry by using CAVER 3.0

| **Tunnel ID** | **Average Bottleneck radius (Å)** | **Priority** | **Bottleneck residues** | **Residues in distance ≤ 3.0 Å from the bottleneck (sorted from the closest)** |
|---|---|---|---|---|
| 1 | 1.087 | 0.57413 | S147 | S161, S146, Q163, K162, E148, I197, E160, V195 |
| 2 | 0.926 | 0.53110 | Q163 | V144, W143, S146, R164, A145 |
| 3 | 0.914 | 0.51491 | W143 | L199, S62, D200, P63, K198, G142 |
| 4 | 0.914 | 0.21400 | W143 | L199, S62, D200, P63, K198, G142 |

**Table S4. R-values for refinements with different possible combinations of degradation products.**

| | A (only **2**) | B (**2**+**4**) | C (**2**+**5**) | D (**2**+**8**+**9**) |
|---|---|---|---|---|
| *R-work* | 0.1258 | **0.1249** | 0.1248 | 0.1253 |
| *R-free* | 0.1468 | **0.1461** | 0.1463 | 0.1457 |

Note: *R-factors* reflect the agreement between the calculated model and experimental x-ray diffraction data. *R-work* and *R-free* are calculated for the "work" and "test" sets, respectively. The test sets include 5% of randomly chosen reflection data. Despite resulting in similar *R-factors*, option B gives a better fit within the electron density map than others and is comparable with SDS analysis and MS results, therefore option B was chosen for final refinement (Fig. S6 - C, D).
